# Use of short-term high energy dietary for estimating transcriptional profiling of liver tissues in sheep

**DOI:** 10.1101/740118

**Authors:** Shaohua Yang, Yunxia Guo, Chengshi He, Yueqin Liu, Yingjie Zhang

**Affiliations:** College of Food and Biological Engineering, Hefei University of Technology, 193 Tunxi Road, Hefei, Anhui 230009, P. R. China; College of Animal Science and Technology, Agricultural University of Hebei, 2596 Lekai South Road, Baoding, Hebei 071001, P. R. China

**Keywords:** RNA-seq, liver metabolism, ewes, DEGs

## Abstract

This study aimed to evaluate if short-term high energy dietary has any stimulatory effects on liver function and metabolic status in sheep. The experiment was carried out using 30 Dorset×Han crossbred ewes (age, 9 ± 0.6 months; weight, 36.58 ± 2.56kg) allocated into two treatments, the control group (DE 11.72 MJ/d; DP 79.71 g/d) and the high energy group (DE18.75 MJ/d; DP 108.44 g/d), respectively. Experiment lasted 20 days, including 10 d for adaption. Blood samples of these ewes were collected to detect the concentrations of glucose, insulin, leptin, and cholesterol, respectively. Then, animals were sacrificed and optimal liver samples subjected to explore the genome-wide transcriptome analysis. Results showed that the weight gain was significantly increased in the high energy group, compared with those in the control group (p< 0.01). The concentrations of glucose, insulin, leptin, and cholesterol were also influenced by short-term nutritional supplementation at different levels. Subsequently, 622 differentially expressed genes were identified by pairwise comparison. Of these, 271 genes were down regulated while 351 genes were up regulated. qRT-PCR analysis of 10 randomly selected genes were consistent with the sequencing results. Gene Ontology (GO) and Kyoto Encyclopedia of Genes and Genomes (KEGG) pathways revealed 12 DEGs (including *PDK4, ABCA9, ALDH6A1, SLC45A3, G0S2, PPARGC1, GHRHR, GHR, DGKI, SOCS2, LPIN1* and *CSKMT*) were significantly enriched in cellular carbohydrate catabolic and metabolic process, phosphorelay sensor and phosphotransferase kinase activity, generation of precursor metabolites and energy, lipid metabolic and transport process, positive regulation of cellular metabolic process, acyl-CoA desaturase activity and monosaccharide metabolic process. Additionally, we concluded an interaction network related to energy metabolism, which might be contributed to elucidate the precise molecular mechanisms of related genes associated with energy metabolism in the liver tissues of sheep.

## INTRODUCTION

Nutrition plays a crucial role in regulating the growth and reproductive performance of farm animals. Energy homeostasis exerts a significant influence on animal health and body condition [1]. Positive energy balance results in increased leptin and insulin concentrations in the blood, which subsequently increased glucose uptake. Previous studies have shown that short-term nutritional supplementation for 4 to 11 days exerts significant influences on the blood metabolites and the reproductive performance of ewes [2–3]. Additionally, it is suggested that nutritional supplementation can directly increase the concentrations of several metabolic hormones in ewes, including glucose, insulin, IGF-1, and leptin [4]. However, to our knowledge, there are few studies on the effect of nutritional supplementation on the liver growth, especially its specific regulatory mechanism. Therefore, the elucidation of the precise molecular mechanisms will have both economic and biological consequences.

As an intermediary between the dietary sources of energy and the extrahepatic tissues, liver is the central organ of metabolism with a considerable energy capacity. In the past several decades, many studies have been conducted to evaluate the effect of feed restriction on liver growth. Feed restriction, which correlates with liver metabolism may exert continuous effects on liver morphology [5] and even induce different degrees of liver impairment [6]. It has also been suggested that maternal undernutrition during late pregnancy could be associated with liver dysfunction [7]. Additionally, nutritional restriction can reduce the liver weights of weaned kids [8] (Yang et al., 2012). Moreover, genomic and proteomic analysis indicates that diet restriction mainly affects liver cell proliferation and apoptosis by the energy-related pathway [9–10]. Despite these findings have contributed significantly to our better understanding of the effect of energy dietary on liver tissues, the precise molecular mechanisms of related genes have not yet to be elucidated.

With the development of high-throughput sequencing technologies, especially RNA-Seq has been widely utilized to explore potential candidate genes which affect important economic traits in animals. In sheep, many studies of RNA-Seq have been conducted using muscle [11], adipose tissues [12], ovarian tissues [13], and abomasal mucosal [14]. However, limited studies on the transcriptome of liver tissues have been reported. The identification of DEGs in liver tissue represents the first step toward clarifying the complex biological properties of metabolic traits. In the present study, we used RNA-Seq technology to examine the genome-wide transcription profile in liver tissues between two groups of sheep with the high energy diet group and the control diet group, respectively. We then proposed key candidate genes affecting liver energy mechanism by conducting integrated analysis. The identified candidate genes could lead to improved selection of sheep while providing new insights into metabolic traits.

## MATERIALS AND METHODS

The experimental design and all animal procedures were authorized by the Institutional Animal Care and Use Committee (IACUC) of Agricultural University of Hebei. In the present study, animals were sacrificed as necessary to ameliorate suffering.

### Animals and samples

The experiment was conducted at Hengshui Shunyao Sheep Farm (Hebei, China; by using 30 Dorset×Han Crossbred ewes (age, 9 ± 0.6 months; weight, 36.58 ± 2.56kg), which were randomly divided into two groups, the control group (DE 11.72 MJ/d; DP 79.71 g/d; n = 15) and the high energy group (DE18.75 MJ/d; DP 108.44 g/d; n= 15), respectively. The diets of the two groups were similar in composition and only varied in energy content, which were outlined in Table 1. The ewes were then placed in individual pens with water and feed available ad libitum.

**Table 1.**
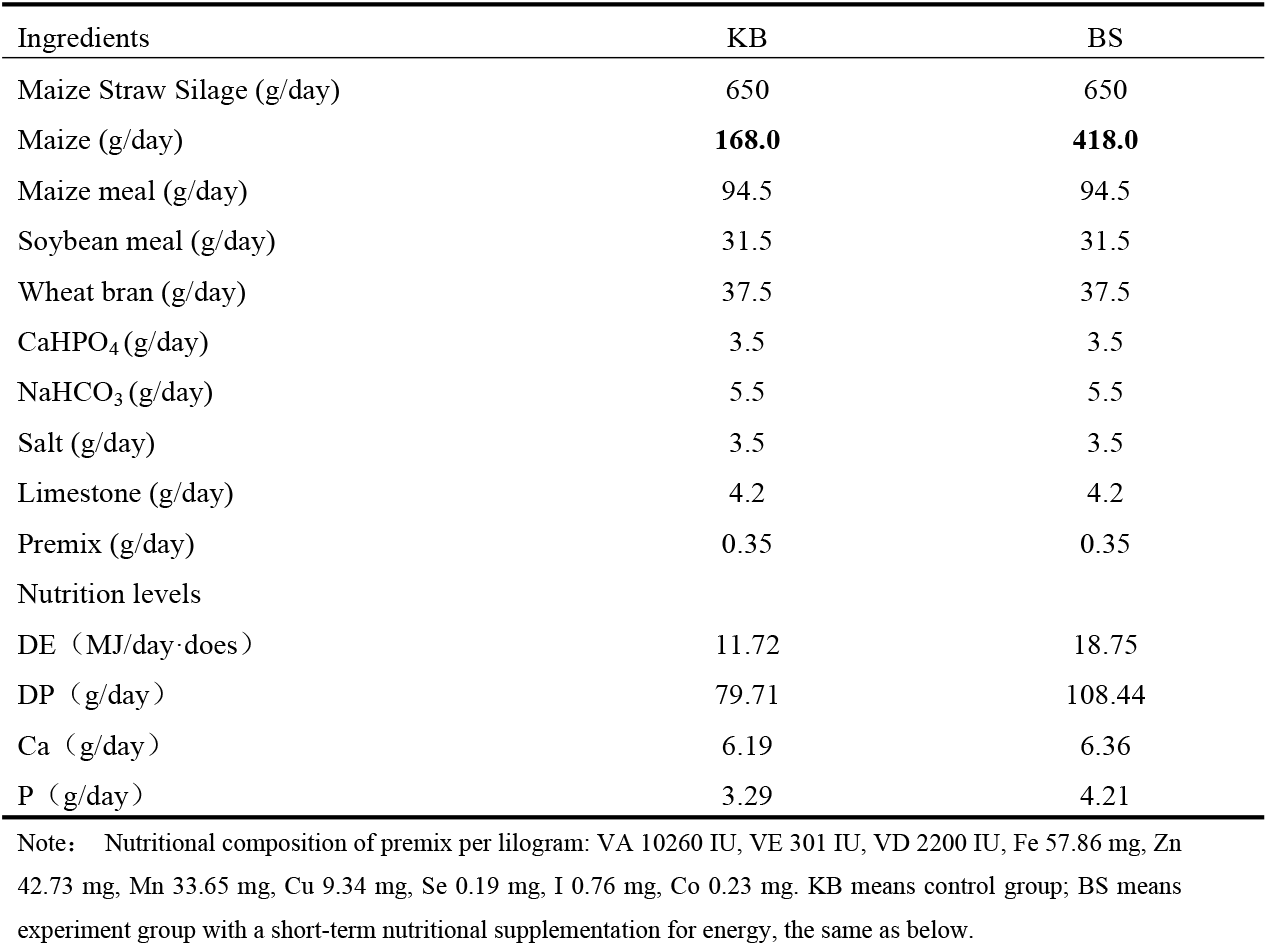
Composition and nutrition levels of TMR

| Ingredients | KB | BS |
| --- | --- | --- |
| Maize Straw Silage (g/day) | 650 | 650 |
| Maize (g/day) | **168.0** | **418.0** |
| Maize meal (g/day) | 94.5 | 94.5 |
| Soybean meal (g/day) | 31.5 | 31.5 |
| Wheat bran (g/day) | 37.5 | 37.5 |
| CaHPO<sub>4</sub> (g/day) | 3.5 | 3.5 |
| NaHCO<sub>3</sub> (g/day) | 5.5 | 5.5 |
| Salt (g/day) | 3.5 | 3.5 |
| Limestone (g/day) | 4.2 | 4.2 |
| Premix (g/day) | 0.35 | 0.35 |
| Nutrition levels |  |  |
| DE (MJ/day·does) | 11.72 | 18.75 |
| DP (g/day) | 79.71 | 108.44 |
| Ca (g/day) | 6.19 | 6.36 |
| P (g/day) | 3.29 | 4.21 |

Note: Nutritional composition of premix per lilogram: VA 10260 IU, VE 301 IU, VD 2200 IU, Fe 57.86 mg, Zn 42.73 mg, Mn 33.65 mg, Cu 9.34 mg, Se 0.19 mg, I 0.76 mg, Co 0.23 mg. KB means control group; BS means experiment group with a short-term nutritional supplementation for energy, the same as below.

Experiment lasted 20 days, including 10 d for adaption. During the adaption period, the animals were fed with the control group diet and energy intake was also strictly monitored. At the end of experiment, these animals were weighed to record the final body weight before slaughtering. Then, blood samples were collected from the jugular vein and centrifuged for 15 min at 3000 × g, then immediately stored at 4°C for subsequent analysis. After slaughtering, liver samples were collected and immediately frozen in liquid nitrogen and stored at −80°C for subsequent analysis.

### Body growth and Tissue glycogen

To evaluate the effect of high energy diet on body growth, the body and liver weights were determined. Meanwhile, tissue glycogen content was measured by colorimetric assay using the Glycogen Assay Kit (Sigma Chemical Co., USA) according to the manufactor’s instructions.

### Glucose, insulin, leptin and cholesterol measurement

The blood samples of these animals during each period were analyzed for glucose (enzymatic colorimetric method using glucose oxidase kit; Huaying Institute of Biotechnology, Beijing, China), insulin and leptin (radioim-munoassay (RIA) procedures, Huaying Institute of Biotechnology, Beijing, China), and cholesterol (cholesterol oxidase method; Jiancheng Institute of Biotechnology, Nanjing, China) in accordance with the instructions of the manufacturer.

### RNA isolation and validation

Total RNA of the liver samples was extracted using the Trizol reagent (Invitrogen, Carlsbad, CA, USA) according to the manufacturer’s instructions. After the quality verification on gel electrophoresis, the concentration and purity of the RNA samples were assessed on a NanoPhotometer^®^ spectrophotometer (Thermo Scientific, Wilmington, DE, USA). The integrity of RNA was assessed using the RNA Nano 6000 Assay Kit of the Bioanalyzer 2100 system (Agilent Technologies, CA, USA).

### RNA sequencing

A total of 3μg RNA from per sample was used as the input material for RNA sample preparations. Based on the manufacturer’s instructions, the transcriptome library was constructed using NEBNext^®^ Ultra™ RNA Library Prep Kit for Illumina^®^ (NEB, USA). Furthermore, TruSeq PE Cluster Kit v3-cBot-HS (Illumina) was used to cluster the index-coded samples on a cBot Cluster Generation System. After cluster generation, the library preparations were sequenced using an Illumina HiSeq 2000 platform, which was followed by FASTQ file generation and the failed reads elimination by CASAVA ver.1.8.2 (Illumina).

### Sequencing data analysis

Using CASAVA ver.1.8.2 (Illumina), the sequencing-derived raw images were transformed into raw reads by base calling. After obtained the raw reads, we removed reads containing low quality reads, adapter and reads containing ploy-N to get clean data through in-house perl scripts. Additionally, the description statistics for the clean data, such as Q20, Q30, and GC-content were calculated for high-quality downstream analysis. The clean data with high quality were used for the downstream analyses.

### Reads mapping

Based on the reference genome, only these reads with a perfect match or one mismatch were further analyzed and annotated. The clean reads were mapped to the reference genome of sheep (Ovis aries version 4.0) using Tophat2 software (version 2.1.0). The index of the reference genome was built using Bowtie v2.2.3 and paired-end clean reads for each individual sample were aligned to the reference genome using TopHat v2.0.12. In addition, HTSeq v0.6.1 was used to count the reads numbers mapped to each gene.

### Differential expression analysis

Differential expression analysis of different groups (the control and high energy groups with different corn content) was identified using the DESeq R package (1.10.1) based on the negative binomial distribution. Furthermore, the Hochberg and Benjamini method was used to adjust the p-values for controlling the false discovery rate [15]. Genes with a FDR value < 0.05 and jlog2-fold changej > 2 were assigned as differentially expressed.

### Functional enrichment analysis

GO and KEGG pathway enrichment analyses of the DEGs were implemented by the Database for Annotation, Visualization and Integrated Discovery (DAVID) website [16] (Huang et al., 2007). GO terms and KEGG pathways with a hypergeometric test from the R package (P < 0.1, FDR-adjusted) were considered significantly enriched among the DEGs. Pathways with fewer than three known genes were discarded.

### Validation of RNA-Seq results

To validate the repeatability and reproducibility of the sequencing results, qRT-PCR was carried out to detect 10 randomly selected DEGs. Primers were designed via Primer3 (http://bioinfo.ut.ee/primer3-0.4.0/primer3/input.htm) and are shown in S1 File. qRT-PCR was carried out in triplicate with the LightCycler^®^ 480 SYBR Green I Master Kit (Roche Applied Science, Penzberg, Germany) in a 15 μ L reaction on a ABI7500 (Applied Biosystems Inc., USA), using the following program: 95°C for 10 min; 40 cycles of 95°C for 10 s, 60°C for 34 s, and 72°C for 10 s; 72°C for 6 min. The mRNA levels of the DEGs were normalized by the housekeeping genes GAPDH and ACTB, and the relative gene expression values were calculated using the 2–ΔΔCt method. Finally, the correlations between RNA-Seq for 10 genes and the mRNA expression level from qRT-PCR were estimated using R (V3.2).

## RESULTS

### Effect of energy level on body and liver weights

To evaluate the effect of energy level on body growth, the body and liver weights were determined, respectively. As shown in Table 2, the item of weight gain was significantly increased in the high energy group, compared with those in the control group (p< 0.01). However, liver weights showed no significant difference between the high energy group and the control group (p> 0.05), which showed that diet had no effect on liver weight.

**Table 2.** Effects on body weight by short-term nutritional supplementation

| Parameter | Initial weight (kg) | Ending weight (kg) | Weight gain(kg) | Liver weight (kg) |
| --- | --- | --- | --- | --- |
| KB | 38.20±2.87 | 38.88±2.78 | 0.68±0.20<sup>A</sup> | 0.56±0.18 |
| BS | 35.87±2.07 | 37.46±2.03 | 1.59±0.21<sup>B</sup> | 0.58±0.19 |

Remark: In the same line, values with different small letter superscripts mean significant difference (P<0.05);
while with different capital letter superscripts mean significant difference (P<0.01).

### Hormone chemical analysis

The concentrations of glucose, insulin, leptin, and cholesterol were influenced by short-term nutritional supplementation at different levels. Also, the highly significant correlation between blood leptin concentrations and other hormones (glucose, insulin, and cholesterol) concentrations was found (p < 0.01), respectively. Meanwhile, as shown in table 3, a highly significant correlation between glucose concentrations and cholesterol concentrations was also detected (p < 0.01) in these ewes.

**Table 3.** Correlation confficients among glucose, insulin, leptin and cholesterol

| Parameters | Glucose (mmol/L) | Insulin (uIU/ml) | Leptin (ng/ml) | Cholesterol(mmol/L) |
| --- | --- | --- | --- | --- |
| Glucose (mmol/L) | 1.000 |  |  |  |
| Insulin (uIU/ml) | 0.253 | 1.000 |  |  |
| Leptin (ng/ml) | 0.405** | 0.452** | 1.000 |  |
| Cholesterol (mmol/L) | -0.301** | 0.066 | -0.472** | 1.000 |

Note: \*\* Indicates extremely significantly correlation (P<0.01); \* Indicates significantly correlation (P<0.05).

Compared with the KB group, higher glucose concentrations were observed for supplemented ewes in the BS group, which ranging from 0.10 to 1.11 mmol/L (mean 0.46 ± 0.36 mmol/L). During the initial (Day 2-4) and final (Day 10) blood, glucose concentrations of the ewes were significantly different (p < 0.05), but not in other days. A significant increase in blood glucose was observed in both KB and BS ewes (figure 1A).

**Figure 1.**
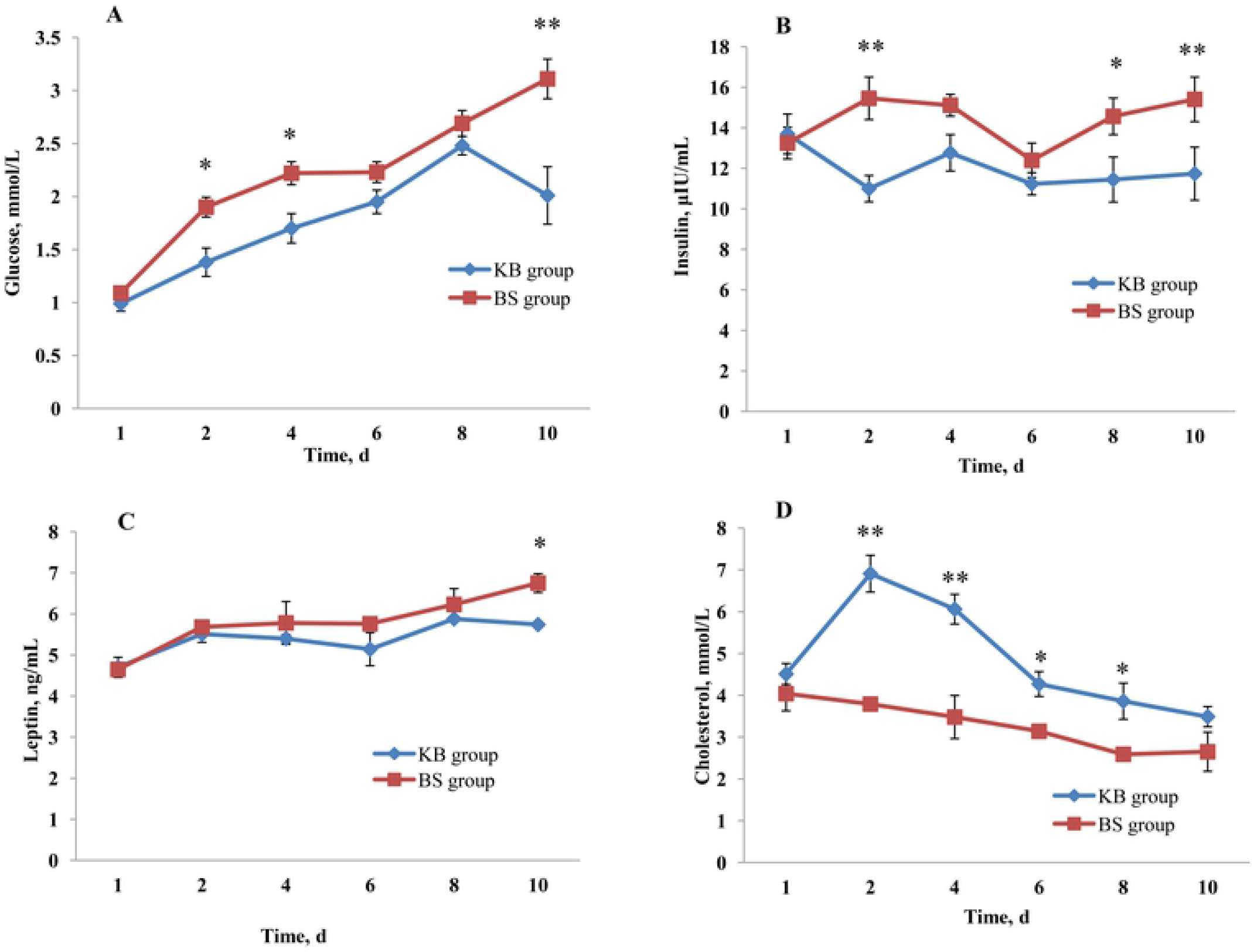
Concentrations of glucose, insulin, leptin and cholesterol in ewes

As shown in figure 1B and figure 1C, in both comparison groups, the insulin and leptin concentrations were higher in BS group. However, during the experimental collection, the ewes had unstable insulin and leptin concentrations. Of these, a highly significant difference (p < 0.01) and significant difference (p < 0.05) were observed between treatments in the insulin concentrations of these ewes at the Day2 and Day8-10, respectively. Additionally, a significant difference (p < 0.05) and highly significant difference (p < 0.05) were also detected between treatments in the leptin concentrations at the Day6 and Day8-10, respectively. However, compared with the KB group, the cholesterol concentration was lower in the BS group (figure 1D), which with a substantially decreasing in blood cholesterol concentration. Moreover, in only the Day2-6 sample collections, significant difference (p < 0.05) were detected in cholesterol concentration.

### RNA sequencing of liver tissue

We acquired a total of 319.20 million clean reads with an average of 53.20 million (range, 47.61 to 57.20 million) for each sample (Table 4). Alignment of the sequence reads against the sheep reference genome (Ovis aries version 4.0) yielded 76.69-80.60% of uniquely aligned reads across the six samples, of which 63.2-73.3% fell in annotated exons, 4.8-8.8% were located in introns, and 18.6-28.0% fell in intergenic regions. The data sets analyzed are available in the NCBI and the SRA ID is SRP151527 (https://submit.ncbi.nlm.nih.gov/subs/sra/SUB4221342/overview). Furthermore, the correlation coefficient (R^2^) between the six individuals was calculated based on the FRPM value of each sample and was shown to be 0.823 −0.956, indicating that the similarity of the three biological samples within each group was sufficiently high (S2 file).

**Table 4.** Basic Statistics of RNA Sequences

| Sample | Raw reads | Clean reads | Error rate(%) | Q20(%) | Q30(%) | GC content(%) |
| --- | --- | --- | --- | --- | --- | --- |
| KB1 | 49659432 | 47614368 | 0.02 | 96.00 | 90.76 | 50.26 |
| KB2 | 55823976 | 53630686 | 0.02 | 95.92 | 90.61 | 51.96 |
| KB3 | 59621642 | 57201632 | 0.02 | 95.94 | 90.67 | 50.25 |
| BS1 | 55194502 | 53105180 | 0.02 | 95.97 | 90.69 | 51.75 |
| BS2 | 57124642 | 54884620 | 0.02 | 96.00 | 90.76 | 51.62 |
| BS3 | 54917556 | 52782450 | 0.02 | 96.21 | 91.17 | 51.18 |

**Note:** Sample name, KB1, KB2, KB3 for the blank group, BS1, BS2, BS3 for the supplementary feed group. Q20, Q30: Calculate the percentage of bases with greater than 20 and 30 Phred values, respectively. GC content: Calculate the percentage of the sum of the number of bases G and C in the total number of bases.

### The identification of DEGs related to energy metabolism

Using the RPKM method, the differential gene expression profile between the control group and the high energy group was examined. In total, 622 genes were detected significantly different between the comparison groups. Of these, 271 genes were down regulated while 351 genes were up regulated. Additionally, the volcanic plot of the two comparison groups was displayed in Figure 2. Furthermore, using integrated analysis of RNA-Seq and gene function, the top 20 genes with the highest absolute value of expression in the liver tissue are shown in Table 5. Interestingly, the energy metabolism associated genes *PDK4, G0S2*, and *CSKMT* accounted for a significant proportion.

**Figure 2.**
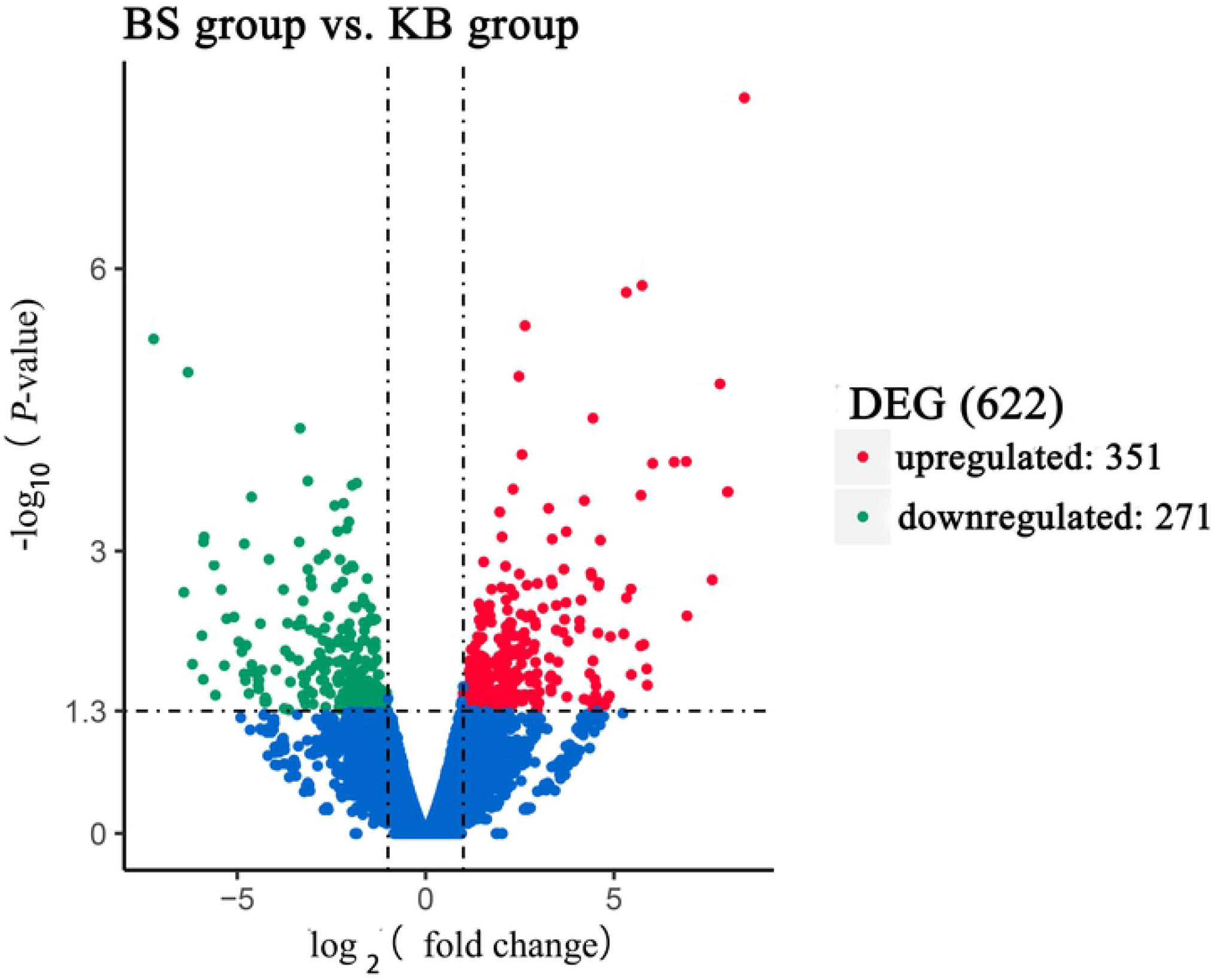
Volcano plot displaying DEGs within two different comparison g

**Table 5.**
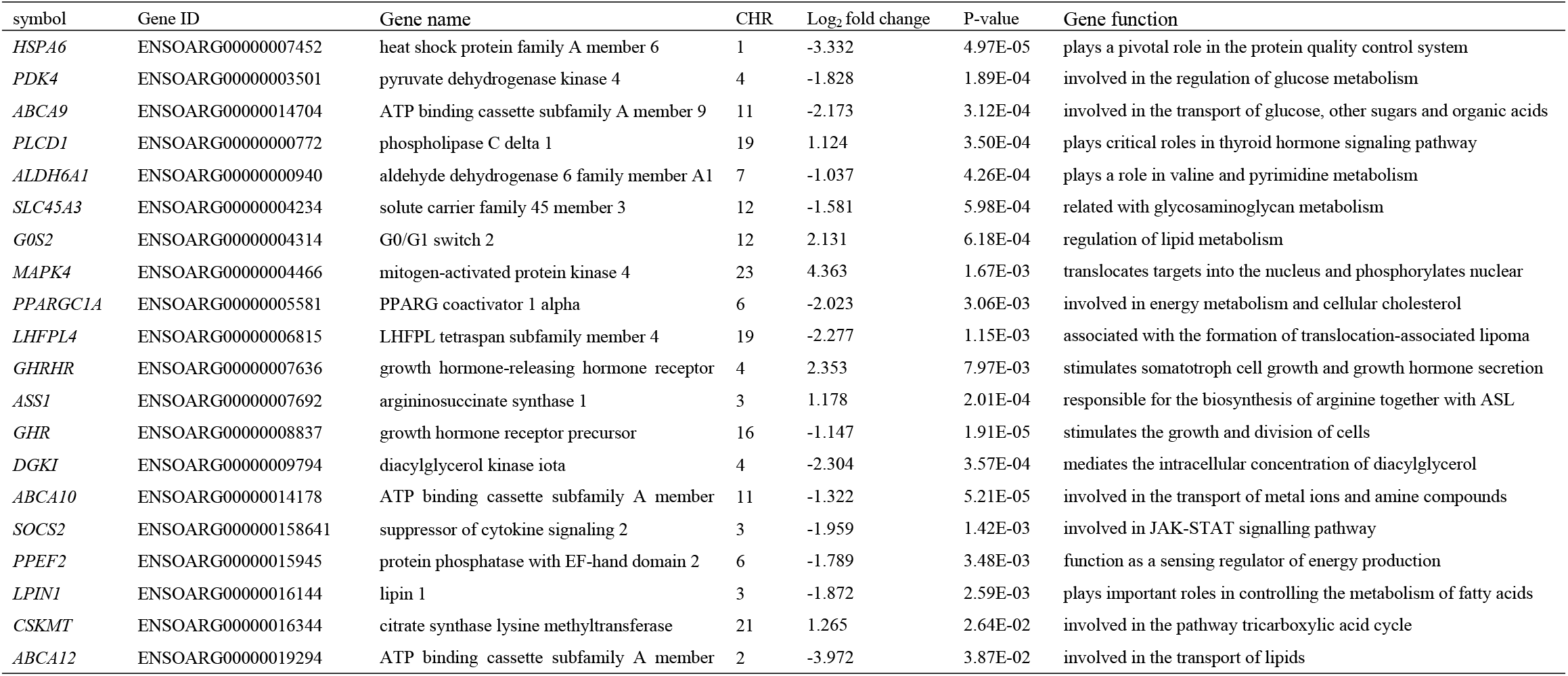
Top 20 differentially expressed genes between high and low IMF content in thigh muscle tissue

### Validation of DEGs

Ten random DEGs (*DQB, PDK4, KATNB1, MBNL3, TPH1, SOCS2, RASGEF1B, FAM210B, WNT2B*, and *NT5C*) were selected for qRT-PCR to validate the RNA-Seq results and the result showed that the correlations between the mRNA expression level of qRT-PCR and RNA-Seq were all consistent (Figure 3). Thus, the reproducibility and repeatability of gene expression data in this study are reliable.

**Figure 3.**
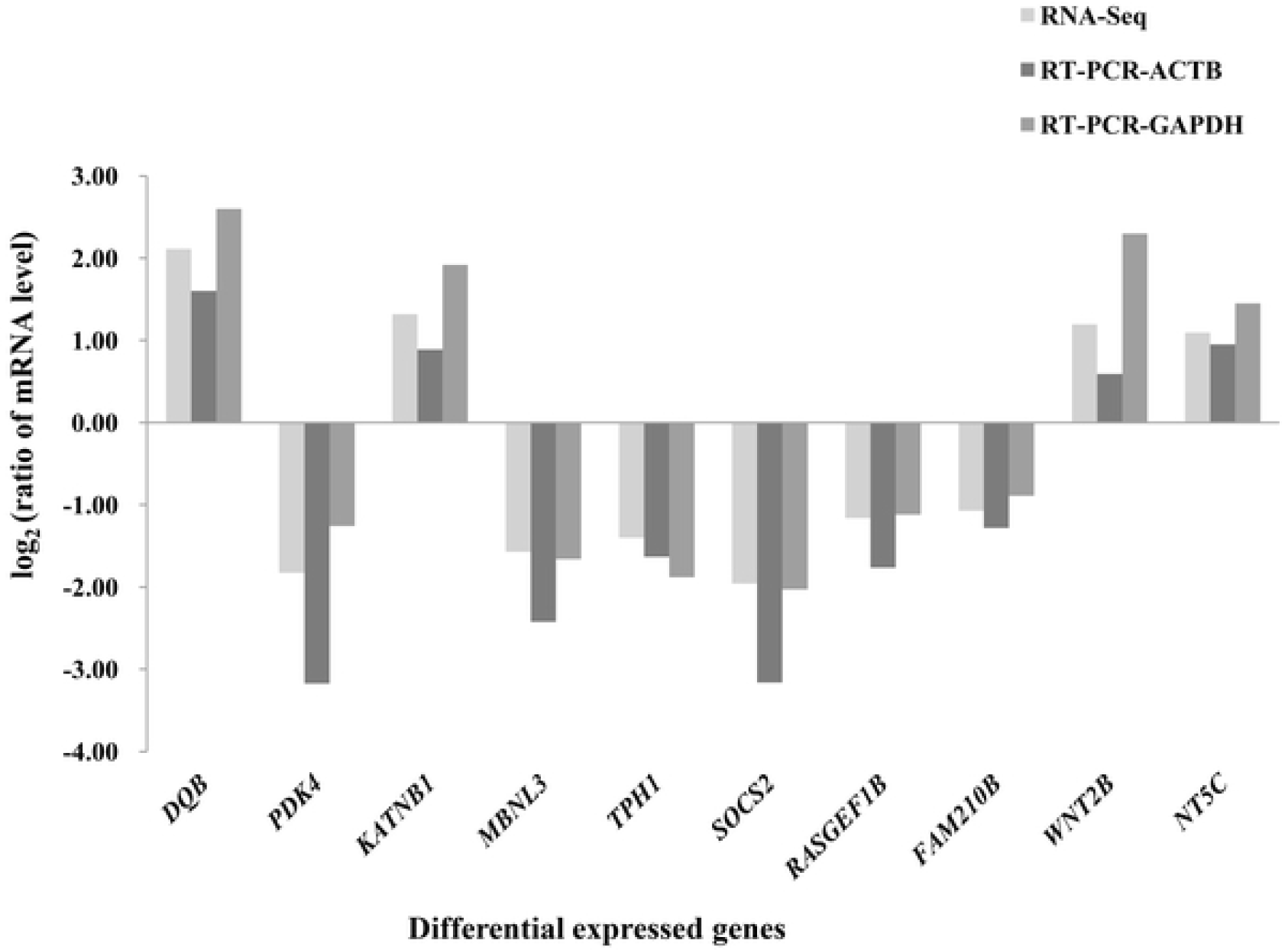
Correlations of mRNA expression level of 10 randomly DEGs bet

### Functional enrichment analysis of DEGs

To gain insight into the biological relationships of genes that differentially expressed in liver tissue between the control group and the high energy group, we performed GO and KEGG pathway enrichment analysis using Database for Annotation, Visualization and Integrated Discovery (DAVID) bioinformatics resource. The results showed that 11 GO biological process terms related to metabolic process of energy and lipid were significantly enriched (P<0.05), which included cellular carbohydrate catabolic process (GO:0044275) and metabolic process (GO:0005975), phosphorelay sensor (GO:0000155) and phosphotransferase kinase activity (GO:0016775), ATPase activity (GO:0016887), generation of precursor metabolites and energy (GO:0006091), positive regulation of cellular metabolic process (GO:0031325), lipid metabolic (GO:0006629) and transport process (GO:0006869), acyl-CoA desaturase activity (GO:0003995) and monosaccharide metabolic process (GO:0005996) (Table 6).

**Table 6.**
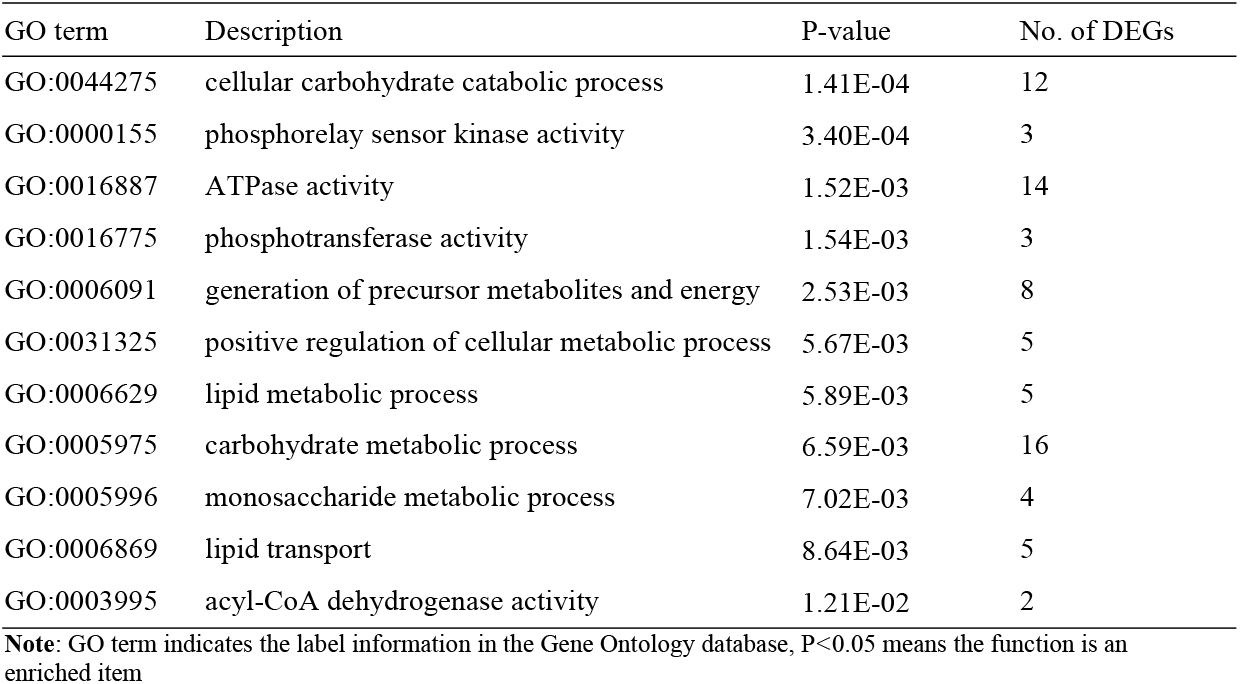
Important GO terms associated with energy metabolism

| GO term | Description | P-value | No. of DEGs |
| --- | --- | --- | --- |
| GO:0044275 | cellular carbohydrate catabolic process | 1.41E-04 | 12 |
| GO:0000155 | phosphorelay sensor kinase activity | 3.40E-04 | 3 |
| GO:0016887 | ATPase activity | 1.52E-03 | 14 |
| GO:0016775 | phosphotransferase activity | 1.54E-03 | 3 |
| GO:0006091 | generation of precursor metabolites and energy | 2.53E-03 | 8 |
| GO:0031325 | positive regulation of cellular metabolic process | 5.67E-03 | 5 |
| GO:0006629 | lipid metabolic process | 5.89E-03 | 5 |
| GO:0005975 | carbohydrate metabolic process | 6.59E-03 | 16 |
| GO:0005996 | monosaccharide metabolic process | 7.02E-03 | 4 |
| GO:0006869 | lipid transport | 8.64E-03 | 5 |
| GO:0003995 | acyl-CoA dehydrogenase activity | 1.21E-02 | 2 |

**Note:** GO term indicates the label information in the Gene Ontology database, $P < 0.05$ means the function is an enriched item

Five KEGG pathways related with glucolipid metabolism were significantly enriched (P < 0.05), including PPAR signaling pathway (oas03320), AMPK signaling pathway (oas04152), glycerophospholipid (oas00564) and glycerolipid metabolism (oas00561) and Insulin signaling pathway (oas04910). As expected, the well-known AMPK signaling pathway (S4 file) and PPAR signaling pathway (S5 file) were found. It is easy to believe that these pathways may be the points for the interaction. Taken together, the proposed molecular regulatory network affecting energy metabolism during liver development is presented in Figure 4.

**Figure 4.**
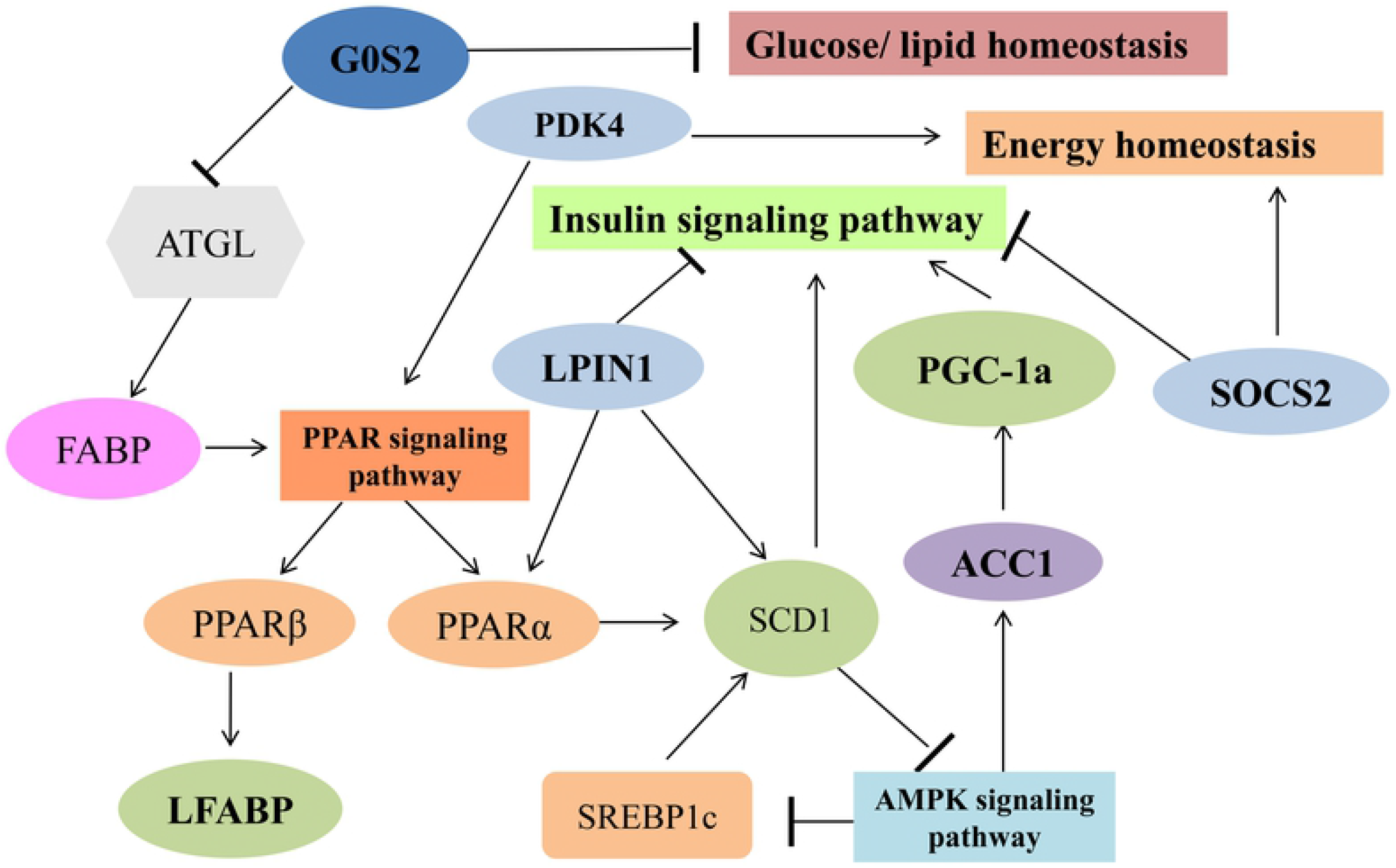
The interaction network of the DEGs associated with energy me

## DISCUSSION

The effect of short-term high energy supplementation on the growth performance, hormones, and liver transcriptome of Dorset×Han crossbred ewes was evaluated in the present study. Herein, maize was selected due to its potential ability to be partially degradable in the rumen of sheep, which provides undegradable starch to flow into the gastrointestinal tract, increasing the entry rate of glucose and other energy substrates into the bloodstream for a longer time [17]. Consequently, the uptake of glucose by the liver may be promoted, stimulating energy metabolism via a direct effect of insulin or insulin-mediated glucose uptake by liver tissues.

The final body and liver weights were increased with a supplementation in energy intake, which agrees with other studies. No significant differences between the dietary treatments were found for ending weight. The lack of differences may possibly be ascribed to the short treatment time and a longer supplementation time was recommended. In addition, the difference in the level of energy between control groups and experimental groups used in this study was possibly not large enough to promote any effect of treatment and a greater concentration of metabolites, which with more times above maintenance requirements, may have a more significant results in ewes [18]. Nonetheless, we still found that the weight gain was significantly increased in the high energy group, compared with those in the control group (p<0.01). These results were expected, as during times of feeding above maintenance level, the usable energy could sufficiently promote the development of young growing animals.

Compared with the does that were not supplemented with maize, mean concentrations of glucose, insulin, and leptin were significantly greater in does supplemented with maize in the present study. This result agrees with some previous studies [19–20], which may be due to diet containing high energy that increased hepatic gluconeogenesis in the rumen. Initially the concentration of glucose rose to a high level that stimulated the secretion of insulin, which showed that the ewes were able to cope with the increasing glucose concentration. Additionally, the highly significant correlation between leptin concentrations and glucose concentrations was found, which may be due to the quantity of nutrients activating ascending pathways in the vagus nerve for independently changing the concentrations of leptin [21–22]. However, an increase in dietary energy leads to reduce plasma total cholesterol. These results were in contrast with previous findings [23–24]. This difference could be attributed to feeding environment, dietary composition, and species-specific differences.

RNA-Seq has many advantages, such as greater dynamic range, removed bias, lower false positives, and higher reproducibility [25]. Furthermore, the correlations between RNA-Seq and the mRNA expression level from qRT-PCR were relatively high. Until now, RNA-Seq has been successfully applied to transcriptome analysis in the sheep [11–14]. In the current study, by comparative analysis and imperative validation, 622 differentially expressed genes were identified by pairwise comparison. Among them, 12 DEGs (including *PDK4, ABCA9, ALDH6A1, SLC45A3, G0S2, PPARGC1, GHRHR, GHR, DGKI, SOCS2, LPIN1* and *CSKMT*) were significantly enriched in energy, glucose and lipid metabolism.

Among them, pyruvate dehydrogenase kinase 4 (*PDK4*) had significantly down-regulated in this study (P < 0.01), which was in accordance with that observed in previous reports. *PDK4*, a member of a family of pyruvate dehydrogenase kinases (PDKs), located in a cluster of similar genes on chromosome 4 and it is highly expressed in the liver, heart and skeletal muscle [4]. In various mammalian tissues, *PDK4* plays a pivotal role in the maintaining energy homeostasis by flexibly attenuating the activity of pyruvate dehydrogenase complex (PDC), which is essential for the catalyzation of pyruvate to acetyl-CoA. *PDK4* deficiency in mice reduced triglyceride (TG) accumulation, which may promote PPARα-mediated fatty acid β-oxidation [26]. Moreover, cumulative evidence has reported that the up-regulation of *PDK4* expression is tightly associated with obesity and diabetes [27]. *PDK4* is a gatekeeper regulator for glucose metabolism and its expression is inhibited by insulin[28], which was consistent with the observed results in our present study.

As a transcriptional coactivator, peroxisome proliferator-activated receptor gamma coactivator 1 alpha (*PPARGC1A* or *PGC-1α*) plays an essential role in metabolic reprogramming in response to dietary availability by controlling the expression of genes involved in glucose and fatty acid metabolism. Some studies showed that the expression of *PGC-1α* in the liver was promoted under the condition of food deprivation, which can stimulate hepatic glycosylation and fatty acid oxidative metabolism [29]. Interestingly, *PGC-1a* knock-out mice display impaired fatty acid oxidation, tricarboxylic acid (TCA) cycle metabolism, and liver gluconeogenesis [30]. On the other hand, Queiroz et al [31] observed an association between *PGC-1a* rs8192678 and higher triacylglycerol and glucose concentration in adolescents, which is similar to the results of an adult study [32]. Similarly, the expression levels of *PGC-1a* had been significantly down-regulated in our study. Thus, we therefore speculated that *PGC-1a* may be a promising candidate gene for liver energy metabolism in sheep.

Lipins (*LPIN1*) are phosphatidate phosphatase (PAP) enzymes, which converts phosphatidic acid (PA) to diacylglycerol (DAG). To date, most studies of mammalian lipin protein function have focused on *LPIN1*. The *LPIN1* gene, encoding lipin1 protein, plays critical roles in lipid homeostasis and metabolism. Recent studies have revealed correlations between *LPIN1* levels and fatty acid metabolism related genes, such as *PPARα, SCD1, PLIN1, ATGL* and *HSL* [33]. In the mouse, *LPIN1* gene mutation can result in lipin-1 deficiency, which causes loss of body fat, fatty liver, and severe insulin resistance [33]. In comparison, over expression of *LPIN1* increases the triglycerides content and systemic insulin sensitivity [34]. Based on these observations in mice, we hypothesized that *LPIN1* is one of the most important candidate genes for energy metabolism in sheep.

Similarly, the expression levels of *GHR, SOCS2, ABCA9, ALDH6A1, SLC45A3*, and *DGKI* had significantly down-regulated in liver tissue (P<0.01). As the previous reported, growth hormone receptor (*GHR*) gene encodes a member of the type I cytokine receptor family involved in the growth and function of many parts of the animals [35]. Suppressor of cytokine signaling 2 (*SOCS2*) is a negative regulator of cytokine signal transduction, which regulates a variety of biological processes, including growth and metabolism, tumorigenesis and immune function [36]. Meanwhile, it is showed that *SOCS2* plays an important role in regulating lipid metabolism by the GH signaling pathway [37]. As a member of the ABC1 subfamily, ATP binding cassette subfamily a member 9 (*ABCA9*) has the important function in the regulation of ATPase activity, transport of glucose and other sugars, and the balance of lipids and cholesterol [38]. Aldehyde Dehydrogenase 6 Family Member A1 (*ALDH6A1*) is highly expressed in the liver and can transform malondialdehyde into acetyl coenzyme A, which is the carbon source of fatty acid synthesis, cholesterol synthesis and ketone formation [39]. Solute carrier family 45 member 3 (*SLC45A3*) are purported to transport sugars, thereby playing an important potential role in maintaining intracellular glucose levels and the synthesis of long-chain fatty acids [40]. However, no previous studies have linked *DGKI* with energy metabolism and further study of these genes seems to be warranted.

On the other hand, the expression levels of *G0S2, GHRHR*, and *CSKMT* had significantly up-regulated in our study. Metabolic regulation is essential for all biological functions. As a multifaceted regulator, the G0/G1 switch gene 2 (*G0S2*) is abundantly expressed in metabolically active tissues and involved in a variety of cellular functions including proliferation, metabolism, apoptosis and inflammation [41]. Particularly, recent studies revealed that *G0S2* acts as a molecular brake on triglyceride (TG) catabolism by selectively inhibiting the activity of rate-limiting lipase adipose triglyceride lipase (ATGL) [42]. Similarly, our study revealed that the expression levels of *G0S2* had significantly up-regulated in liver. Growth hormone releasing hormone receptor (*GHRHR*) is a membrane associated receptor, which can affect cell proliferation and growth hormone secretion of animals by increasing intracellular cAMP. However, the precise biological functions of *GHRHR* and *CSKMT* are not known and further research is required to understand the molecular mechanisms of these genes on energy metabolism in sheep.

Meanwhile, the regulatory network underlying energy metabolism of livers was explored by KEGG pathway analysis in sheep. As expected, the well-known AMPK signaling pathway and PPAR signaling pathway were found. Of special interest, three pathways (glycerophospholipid and glycerolipid metabolism and Insulin signaling pathway) also were enriched, and it was revealed that these pathways may be the points for the interaction. These findings provide new clues for revealing the molecular mechanisms underlying energy metabolism of livers in sheep. This novel speculation and its detailed mechanism through pathways related to energy metabolism identified here needs further validated.

### Conclusions

The study provided a comprehensive point of the complexity of the liver tissues, and revealed 622 differentially expressed genes between the control group (DE 11.72 MJ/d; DP 79.71 g/d) and the high energy group (DE18.75 MJ/d; DP 108.44 g/d). Comprehensive analysis of differential gene expression, biological functions, GO enrichment and pathway analysis showed that 12 DEGs (including *PDK4, ABCA9, ALDH6A1, SLC45A3, G0S2, PPARGC1, GHRHR, GHR, DGKI, SOCS2, LPIN1* and *CSKMT*) were significantly enriched in cellular carbohydrate catabolic and metabolic process, phosphorelay sensor and phosphotransferase kinase activity, generation of precursor metabolites and energy, lipid metabolic and transport process, positive regulation of cellular metabolic process, acyl-CoA desaturase activity and monosaccharide metabolic process. Additionally, we concluded an interaction network related to energy metabolism, which might be contributed to elucidate the precise molecular mechanisms of related genes associated with energy metabolism in the liver tissues of sheep.

## Supporting information

S1 File. PCR primers for qRT-PCR validation of 10 DEGs between the two different comparison groups. (PDF)

S2 File. The distribution of reads in different regions of the reference genome. (PDF)

S3 File. Correlation between biological replicates within three samples. (PDF)

S4 File. AMPK signaling pathway. (PNG)

S5 File. PPAR signaling pathway. (PNG)

## Acknowledgments

This work was supported by the National Key R&D Program of China (2018YFD0502100) and National Science Foundation Project of Anhui (1908085MC63).

